# Recency order judgments in short term memory: Replication and extension of Hacker (1980)

**DOI:** 10.1101/144733

**Authors:** Inder Singh, Marc W. Howard

## Abstract

The classic finding from short-term relative JOR tasks is that correct response time (RT) depends on the lag to the more recent item but not to the less recent item (Hacker, 1980). For decades, researchers have argued that this finding is consistent with a self-terminating backward scanning model (Muter, 1979; Hacker, 1980; Hockley, 1984; McElree & Dosher, 1993). This finding has taken on new importance in light of recent proposal that many forms of memory depend on a compressed representation of the past (Howard, Shankar, Aue, & Criss, 2015). This paper replicates and extends the results of the classic papers. A Bayesian t-test showed substantial evidence for the null effect of lag to the less recent item on correct RT. In addition, this paper reports that correct RT is a sub-linear function of lag to the more recent probe and replicates the classic finding that error RT depends on lag to the less recent probe. These findings place new constraints on models of short-term memory scanning.

---

In a relative judgment of recency (JOR) task participants choose which of two probes from a list was presented more recently. Recency in this task is traditionally measured in units of lag, which is the number of time steps in the past at which the probe was presented. That is, if the last item in the list was presented as a probe, it would be associated with a lag of one. The classic finding from short-term relative JOR tasks is that correct response time (RT) depends on the lag to the more recent item but not to the less recent item (see Fig. 1, Muter, 1979; Hacker, 1980; Hockley, 1984). For decades, researchers have argued that this finding is consistent with a self-terminating backward scan (Muter, 1979; Hacker, 1980; Hockley, 1984; McElree & Dosher, 1993). Because the scan starts at the present and proceeds backwards in time, RTs show a recency effect. Because the scan is self-terminating this naturally accounts for the finding that correct RTs do not depend on the lag of the less-recent probe. This account also naturally explains the finding that incorrect RTs depend on the lag to the (incorrectly) selected probe item.

**Figure 1.**
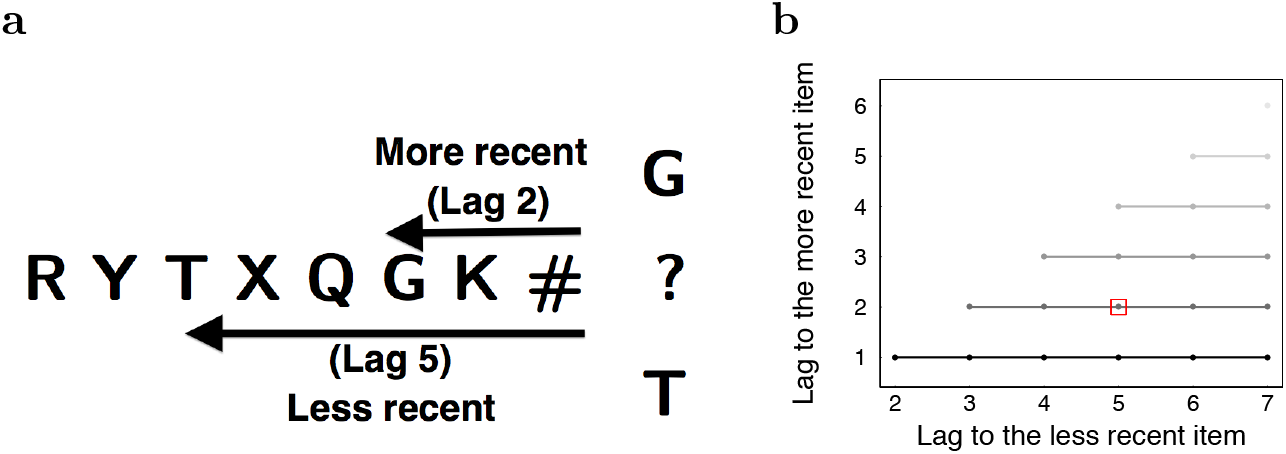
Schematic of the judgment of recency (JOR) task. **a**. The participants are shown a list of letters followed by a probe containing two letters from the list. The participants are required to choose the more recent of the two probe items. **b**. The lag combinations in this task. The most recent lag, lag 1 can be paired with a less recent probe from six possible lags, lags 2 through 6. This is represented by the darkest line. The shading represents the lag to the more recent item and the less recent item is plotted on the x-axis. The red box indicates the probed lag in the schematic on the left. We follow this shading convention in the subsequent plots where both the less and the more recent lags are included on the same graph.

A backward scanning model implies that memory for the list is organized along a temporal axis; in order for memory to be sequentially “scanned” it must be organized in some way. Recently, this finding has taken on new theoretical importance, with some authors arguing that many different forms of memory could be supported by different operations on a temporally-organized memory (Howard et al., 2015). This places additional explanatory importance on the null effect of lag to the less recent probe. Moreover, this theoretical proposal argues that the temporal dimension should be compressed, consistent with longstanding arguments about memory (Brown, Neath, & Chater, 2007; Balsam & Gallistel, 2009). A serial scan of a compressed temporal representation should result in a sublinear increase in RTs, rather than a linear increase in RTs as one would expect from scanning of an uncompressed representation of time. Although previous results are consistent with a sublinear increase in RT with lag (see e.g., Fig. 7 of Hockley, 1984, Fig. 5 Hacker, 1980) this was not statistically evaluated in those prior studies.

This study describes findings from a short-term JOR task very similar to the procedure used in (Hacker, 1980). To anticipate the results, the findings support the results of the classic papers with modern statistics—we will argue that correct response time does not depend on the lag to the less-recent item as evaluated using a Bayesian t-test. In addition, we found that the response time is a sub-linear function of lag to the more recent item. This finding is as predicted by scanning of a compressed representation. If performance in the JOR task results from scanning of a temporal representation of the past, the sublinear increase in RT suggests that this representation is compressed along the temporal axis.

## Method

### Procedure

The procedure of this experiment follows the procedure of Experiment 2 of Hacker (1980) closely. We describe the procedures in detail here, noting differences between this experiment and prior work where relevant. Participants were presented with a list of 9, 11, or 13 consonants at the rate of 5.5 letters per second. At the end of the list, two of the last seven letters were chosen randomly and the participants were asked to indicate using left or right arrow key which of the two letters had appeared more recently. In Figure 1, G and T are presented as the probe items. Because G was presented more recently than T, the correct answer is G. In addition, participants were asked to respond with the up arrow key if they did not remember seeing either of the probe letters on the list. If the participant did not make a response within 6 s, the trial was terminated. Less than .004 of trials terminated without a response.

The distance to the more recent probe stimulus was varied from lag 1 (the last stimulus in the list) to 6. The lag to the less recent probe was varied from 2 to 7. This leads to 21 possible lag combinations, which were presented in a random order. Each participant completed 320 trials.

There are several methodological differences between this procedure and the procedure of Experiment 2 of Hacker (1980). Unlike the Hacker (1980) study, in this experiment participants were never given foils that did not appear in the list. Also, in the Hacker (1980) study participants were not given the option to respond indicating that they did not remember either of the probes. The participants in the Hacker (1980) study were also more experienced in the task, experiencing a variety of presentation rates over several experimental sessions. In this experiment, participants received only one presentation rate in one session lasting about forty minutes.

### Participants

The participants that participated in the study were drawn from the participant pool for Boston University’s introductory psychology class. The study materials and protocol was approved by the Institutional Review Board at Boston University. 108 participants signed up for the study. One participant withdrew from the experiment. Data from 11 participants was excluded because their overall accuracy across all lags was no better than chance. Data from the remaining 96 participants was analyzed using R.

## Results

On .17 ± .02 of trials, participants selected neither of the probes, which we refer to here as “abstain” responses. The overall median RT for the abstain responses was 1.5 ± .2*s*. This was slower than the median response time in the slowest condition (lag 6) for both the correct and incorrect choices (*t*(187) = −2.2, *p* < 0.05). These abstain trials are excluded from the subsequent analyses. The accuracy and response time calculations that follow are calculated based on the trials on which the participant made a choice.

### Strategy for analyzing the two lag variables

The analyses that follow consider the effect of two variables, lag to the more recent probe and lag to the less recent probe. In many cases, the effect of one or more of these variables is clearly non-linear. Simply putting both lags into a linear model under these circumstances would raise the possibility that a spurious effect of one lag would result from an attempt to account for residuals of the other lag. Moreover, because of the definition of lag, the values of the two lag variables covary (note that the lines in Fig. 1b are of different lengths). The probe at the more recent lag was compared to the probe at lags further in in the past. For each more recent lag, there is a variable number of less recent lags. In order to control for the effect of unequal number of combinations, we adopt a two stage analysis strategy.

First, we take the less salient of the two variables and compute a distance effect for that variable. If that distance effect is different from zero (as assessed by a Bayesian t-test), we allow that variable to enter into a linear mixed effect analysis. In cases where the distance effect is not reliable, the function of the linear mixed effect analysis is simply to determine whether the apparent effect of the more salient variable is statistically reliable.

To be more concrete, faced with data that looks like Figure 1b, we would, for each value of the more recent lag (the different lines in Fig. 1b), compute the slope with respect to the lag to the less recent probe for each participant. If we found evidence that the distance effect was zero (as assessed with a Bayesian t-test (Rouder, Speckman, Sun, Morey, & Iverson, 2009)) we would not include it as a factor in a linear mixed effect analysis. The purpose of this analysis would simply be to confirm that there is an effect of the lag to the more recent probe. This strategy applies to both analyses of accuracy and correct RTs. In the case of error RTs, the salience of the two lags is reversed, consistent with prior results (Hacker, 1980).

### Accuracy depended on both the lag to the more recent probe and the lag to the less recent probe

The probability that participants selected the more recent probe was .70 ± 0.01. The accuracy was .82 ± .01 when the lag of the more recent probe was 1 and dropped to .49 ± .02 when the lag to the more recent lag was six. At lag 6 the probability of choosing the more recent probe was not different from chance (Chi-squared prop test, *χ*^2^(96) = 89.1, p-value not significant). Lag 5 had an accuracy of 0.56±0.1 and was significantly higher than chance (Chi-squared prop test, *χ*^2^(96) = 142.6, *p* < 0.01). Accuracy monotonically increased at lower lags and was also significantly higher than chance.

Accuracy also depended on both the lag to the more recent probe and the lag to the less recent probe. For a given lag to the more recent probe, the accuracy improved as the lag to the less recent probe increased (distance effect). The upward-sloping lines in Figure 2a indicate the presence of a distance effect. To quantify the distance effect for each participant, we calculated the slope of each line in Figure 2a. A Bayesian t-test (Rouder et al., 2009) on the obtained slopes revealed “decisive evidence” (Wetzels & Wagenmakers, 2012; Kass & Raftery, 1995; Jeffreys, 1998) favoring the hypothesis that the slopes are different from 0 (JZS Bayes Factor > 10^2^).

**Figure 2.**
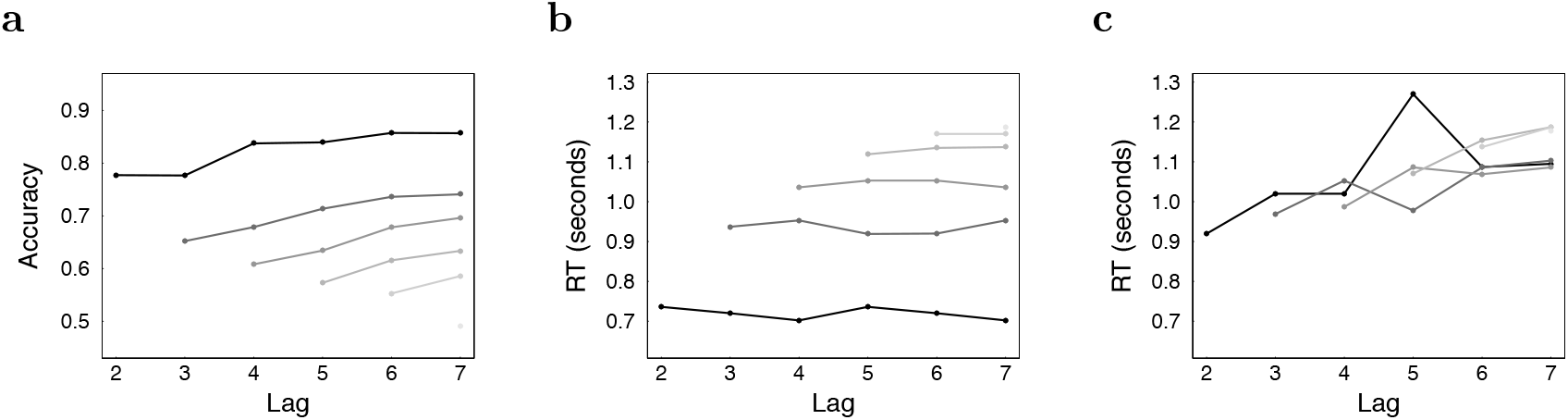
Accuracy, correct RT and incorrect RT are plotted as a function of the lag to the less recent probe. Different lines represent different values of the lag to the more recent probe. (darker lines correspond to more recent lags). **a**. Accuracy depends on the lag to the more recent item and also shows a weak distance effect (note that the lines are not flat). **b**. Correct RT depends strongly on the lag to the more recent probe. The flat lines suggest that there was not an effect of the lag to the less recent probe (see text for details). **c**. Incorrect RT for incorrect responses depends on the lag to the less recent probe, but at most weakly on the lag to the more recent probe (see text for details).

The effects of the two lags on accuracy was quantified using a linear mixed analysis with independent intercepts for each participant. The accuracy decreased with an increase in the lag to the more recent probe by .078 ± .002, *t*(1918) = −31.9, *p* < 0.01 per unit change in lag. Accuracy also increased with the lag to the less recent probe by .023 ± .002, *t*(1918) = 9.73, *p* < 0.01 per unit change in the lag. These findings are consistent with the findings from prior studies.

### Correct response time depended strongly on the lag to the more recent probe but not on the lag to the less recent probe

The response times for the correct responses depended strongly on the more recent lag as seen in Figure 2b. The median response time varied from .72±.02 s for the most recent lag to 1.36 ± .06 s for a lag of six. In contrast to the distance effect seen in accuracy Figure 2a, the lines in Figure 2b appear to be flat. In order to assess this distance effect more directly, we calculated the slopes of lines in Figure 2b separately for each participant and performed a Bayesian t-test (Rouder et al., 2009) on the slopes. This analysis showed “substantial evidence” (Wetzels & Wagenmakers, 2012; Kass & Raftery, 1995; Jeffreys, 1998) favoring the hypothesis that the slopes are not different from 0 (JZS Bayes Factor = 3.3). A linear mixed effects analysis allowing for independent intercepts for each participant showed a significant effect of the lag to the more recent probe, .124 ± .006 s, *t*(478) = 21.6, *p* < 0.001. These results replicate prior studies, but extend them by establishing positive evidence for the null using the Bayesian t-test.

### Response time varies sub-linearly by lag to the more recent item

Figure 2b suggests that correct RTs depended prominently on the lag to the more recent item. Further it appears that the spacing between these lines goes down as the lag increases. This suggests that the RT depends sub-linearly on the lag to the more recent probe, as predicted by a backward self-terminating scanning model that scans along a temporally-compressed representation.

In order to evaluate whether scanning times depended on lag to the more recent probe in a sublinear fashion, we compared two models. In one model, RT was regressed onto the more recent lag. In the other RT was regressed onto the logarithm of the more recent lag. The log model fit better than the linear model, Δ*AIC* = 4.1 (log model is 60.34 times more likely as compared to the linear model).

As an additional test for sublinearity we also compared the linear model to polynomial models with various powers of lag to the more recent probe. A quadratic model fit better than the linear model Δ*AIC* = 3.7. The quadratic term had a negative regression coefficient −0.6 ± 0.2, *t*(477) = −2.6, *p* < 0.01. Including higher order terms did not further improve the fit. Consistent with the conclusions of the logarithmic analysis reported above, both approaches found evidence that the effect the lag to the more recent probe on correct RT was sublinear.

The finding of sublinearity is visually consistent with prior studies (Hacker, 1980; Hockley, 1984). However to our knowledge it had not previously been statistically evaluated.

### Incorrect response time depended strongly on the lag to the less recent probe

In a self-terminating backward scanning model, if the scan misses the more recent probe, it would then terminate on the less recent probe. These responses would be errors and the scanning time for these errors would depend on the lag to the less recent probe.

Given the overall error rate of .30 ± 0.01, there are less than half the number of observations for incorrect RTs as there are for correct RTs. Also note that the number of errors is not evenly distributed over lags, so that some points in Figure 2c have many fewer observations than others. Nonetheless, error RTs appear to depend reliably on the lag to the less recent item (note that all the lines increase from left to right). There does not appear to be a strong effect of the lag to the more recent probe (note that the different lines tend to lie on top of one another).

To evaluate whether there was an effect of the lag to the more recent probe we calculated the slope of the distance effect for each value of the lag to the less recent probe. This is analogous to the distance effect calculation for correct RTs except the distance effect is calculated separately for the lag to the less recent probe rather than the more recent probe. That is, for errors we computed a slope for each cluster of points in Figure 2c rather than across each line. A Bayesian t-test showed “strong evidence” (Wetzels & Wagenmakers, 2012; Kass & Raftery, 1995; Jeffreys, 1998) favoring the hypothesis that the slopes of the median response times as a function of the more recent lag are not different from 0 (JZS Bayes Factor = 14.5).

A linear mixed effects analysis, allowing each participant to have an independent intercept, and the less recent lag as regressor showed a significant effect of the lag to the less recent probe on the median response time of incorrect responses, .033 ± .007 s, *t*(473) = 4.9, *p* < 0.001.

## Discussion

This study replicated the classical finding in a relative JOR task that correct RT depends on the lag to the more recent probe but not on the lag to the less recent probe (Fig. 2b). Beyond simply failing to observe a positive effect, a Bayesian t-test found substantial evidence that there was no effect of the less recent probe on correct RT. In contrast to the findings for correct RTs, the pattern was reversed for error RTs. For error RTs, the lag to the less recent probe showed a robust effect, while a Bayesian t-test found strong evidence that there was no effect of lag to the more recent probe (Fig. 2c). For both correct and error RTs, the lag to the probe stimulus that was selected had a large effect on RT whereas the lag to the probe that was not selected did not have an effect on RT.

Scanning models posit that order information for the items in memory drives access, such as in conveyor belt models of memory (Murdock, 1974). The dependence of RT on only the lag of the selected stimulus is a strong prediction of a self-terminating scanning model. In contrast, strength-based models predict a distance effect on RTs in a relative JOR task, i.e. RT should depend on both lags. Although there is not a distance effect on correct RT, there is a distance effect on accuracy. Scanning models have addressed this by assuming that the probability of the search terminating when a probe is encountered goes down with increasing lag (Hacker, 1980; Howard et al., 2015).

This paper also showed that the effect of lag to the more recent probe on correct RT was sublinear (Fig. 3). This is a prediction of the hypothesis of a model in which the self-terminating search scans over a compressed temporal representaiton (Howard et al., 2015). This result is also consistent with other conceptions of the scanning process as well. For instance the model presented in Hacker (1980) assumes that list stimuli are associated with an availability that falls off with increasing lag. The amount of time to scan a list stimulus is a function of its availability. Because availability decreases, the scanning rate appears to accelerate as one proceeds towards the past.

**Figure 3.**
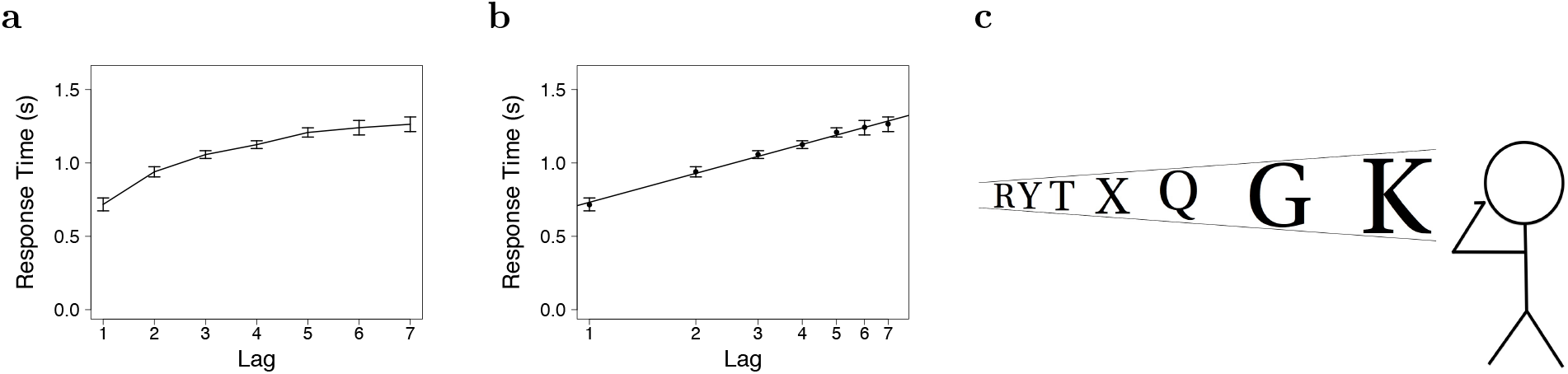
Response time varies sub-linearly with lag. **a**. Median RT plotted as a function of the lag to the chosen item. In case of correct responses this is the lag to the more recent item and in case of errors this is the lag to the less recent item. **b**. Median RT plotted as a function of the log of the lag to the chosen item. The median response times are well described by a straight line on a log scale. **c**. Schematic of the underlying memory representation inferred from the results obtained in the experiment. The accuracy goes down with the lag to the more recent item and this is represented as change in the strength (smaller sized letters). The spacing between the lines is foreshortened to indicate compression in the underlying memory representations.

### Is scanning under strategic control?

A scanning model implies that the brain maintains some representation organized such that attention can be sequentially deployed along a temporal dimension. This naturally leads to the question of whether that deployment of attention is under strategic control or not. The scanning rate one would infer from the present results is such that one can scan a few seconds into the past in about one second of search time. The relative slowness of this putative scan relative to other cognitive processes suggests it could be under strategic control.

Additional evidence suggesting that scanning is under strategic control comes from a variant on the relative JOR task in which the instructions are reversed. Chan, Ross, Earle, and Caplan (2009) had participants perform either a JOR task or a relative order judgment in which they were asked to select the probe stimulus that appeared *earlier* in the list. Unsurprisingly, their JOR task replicated the canonical results from relative JOR, with correct RTs failing to show an effect of the lag to the less recent probe, consistent with a backward self-terminating scanning model. Remarkably, however, when the instructions were reversed the pattern of results for correct RTs was consistent with a *forward* self-terminating scanning model (see also Liu, Chan, & Caplan, 2014). This suggests that a temporally-organized representation can be accessed strategically.

### Relationship to other JOR tasks over longer time scales

In addition to the short-term relative JOR task studied here, investigators have studied recency judgments using a variety of similar methodologies. Yntema and Trask (1963) first introduced the relative JOR task as a continuous judgment in which a stream of stimuli was occasionally interupted by a pair of probes. They examined a much wider range of lags than in the short lists used here. The standard finding from continuous JORs is that accuracy decreases, all else equal, as the distance between the two probes decreases and as the lag to the more recent probe increases. These findings are broadly consistent with the changes in accuracy observed in relative JOR on short lists of stimuli (such as this study).

In studies of absolute JOR, participants rate the distance to the probe numerically (Hinrichs & Buschke, 1968; Hinrichs, 1970; Hintzman, 2010; Hacker, 1980). The canonical findings from absolute JORs are 1) that the variability in ratings increases for probes further in the past and 2) participants’ ratings for the lag of a probe decreases sublinearly with the actual lag of the probe. (Hinrichs & Buschke, 1968) argued that a logarithmic function relates rated lag to actual lag. Notably, when a probe stimulus was presented multiple times in the past, participants can rate the separate occurrences nearly independently. Hintzman (2010) presented stimuli three times, P1, P2 and P3. Participants rated the recency of the most recent presentation at both P2 and at P3. The key finding was that ratings at P3 depended a great deal on the lag between P2 and P3, but very little on the lag between P1 and P2 (see also Flexser & Bower, 1974).

### A common model for JORs across scales?

While none of the findings discussed here is sufficient in isolation to argue for a memory representation that utilizes temporal ordering (Friedman, 1993), it is worth noting that all of these findings can be explained using the same model (Howard et al., 2015). Suppose first that all of the tasks described rely on a logarithmically-compressed timeline. In this representation, past experience is represented along a timeline. Consistent with the Fechner law, the time axis is logarithmically compressed, resulting in a scale-invariant representation (Chater & Brown, 2008; Brown et al., 2007). This property allows behavioral models to exhibit similar properties over time scales ranging from a few seconds (e.g., this study Hacker, 1980) up to many minutes (e.g., Yntema & Trask, 1963). A backward self-terminating scan over a timeline with these properties would naturally lead to the results in this paper, including the sublinear function relating correct RT to the lag to the more recent probe (Fig. 3).

However, the results from continuous JOR and absolute JORs over longer time scales are also at least roughly consistent with this model (for detailed models see Howard et al., 2015). For instance, the logarithmic function relating judged lag to actual lag is a natural consequence of logarithmic compression. Similarly, the minimal dependence of judged recency on previous presentations of a stimulus (Hintzman, 2010) is a natural consequence of a backward self-terminating model. The forward scanning results (Chan et al., 2009; Liu et al., 2014) could result from recovery of the temporal context at the start of the list (Davelaar, Goshen-Gottstein, Ashkenazi, Haarmann, & Usher, 2005) followed by a forward scan using a translation operator (Shankar, Singh, & Howard, 2016). The major gap in reconciling short-term (this study, Muter, 1979; Hacker, 1980; Hockley, 1984; McElree & Dosher, 1993) and long-term (Hinrichs & Buschke, 1968; Hintzman, 2010; Yntema & Trask, 1963) JORs is the lack of RT data for judgments over scales more than a few seconds.

## Conclusions

This study examined response times and accuracy in short-term JOR tasks. We replicated and extended classic results from the short-term JOR task. Whereas accuracy showed a distance effect, both correct and error RTs depended only on the lag to the probe stimulus that was chosen. Bayesian analyses showed positive evidence for a null effect of the lag to the probe that was not chosen. Moreover, the increase in correct RT was a sublinear function of the lag to the more recent probe. These findings are consistent with a backward self-terminating scanning model in which the participant scans over a logarithmically-compressed representation of the past.

## References

Balsam, P. D., & Gallistel, C. R. (2009). Temporal maps and informativeness in associative learning. Trends in Neuroscience, 32(2), 73–78.

Brown, G. D. A., Neath, I., & Chater, N. (2007). A temporal ratio model of memory. Psychological Review, 114 (3), 539–76.

Chan, M., Ross, B., Earle, G., & Caplan, J. B. (2009). Precise instructions determine participants’ memory search strategy in judgments of relative order in short lists. Psychonomic Bulletin & Review, 16(5), 945–51.

Chater, N., & Brown, G. D. A. (2008). From universal laws of cognition to specific cognitive models. Cognitive Science, 32(1), 36–67. doi: 10.1080/03640210701801941

Davelaar, E. J., Goshen-Gottstein, Y., Ashkenazi, A., Haarmann, H. J., & Usher, M. (2005). The demise of short-term memory revisited: empirical and computational investigations of recency effects. Psychological Review, 112(1), 3–42.

Flexser, A. J., & Bower, G. H. (1974). How frequency affects recency judgments: a model for recency discrimination. Journal of Experimental Psychology, 103(4), 706–16.

Friedman, W. J. (1993). Memory for the time of past events. Psychological bulletin, 113(1), 44.

Hacker, M. J. (1980). Speed and accuracy of recency judgments for events in short-term memory. Journal of Experimental Psychology: Human Learning and Memory, 15, 846–858.

Hinrichs, J. V. (1970). A two-process memory-strength theory for judgment of recency. Psychological Review, 77(3), 223–233.

Hinrichs, J. V., & Buschke, H. (1968). Judgment of recency under steady-state conditions. Journal of Experimental Psychology, 78(4), 574–579.

Hintzman, D. L. (2010). How does repetition affect memory? Evidence from judgments of recency. Memory & Cognition, 38(1), 102–15.

Hockley, W. E. (1984). Analysis of response time distributions in the study of cognitive processes. Journal of Experimental Psychology: Learning, Memory, and Cognition, 10(4), 598–615.

Howard, M. W., Shankar, K. H., Aue, W. R., & Criss, A. H. (2015). A distributed representation of internal time. Psychological review, 122(1), 24.

Jeffreys, H. (1998). The theory of probability. OUP Oxford.

Kass, R. E., & Raftery, A. E. (1995). Bayes factors. Journal of the American Statistical Association, 90, 773–795.

Liu, Y. S., Chan, M., & Caplan, J. B. (2014). Generality of a congruity effect in judgements of relative order. Memory & Cognition, 1–20.

McElree, B., & Dosher, B. A. (1993). Serial recovery processes in the recovery of order information. Journal of Experimental Psychology: General, 122, 291–315.

Murdock, B. B. (1974). Human memory: Theory and data. Potomac, MD: Erlbaum.

Muter, P. (1979). Response latencies in discriminations of recency. Journal of Experimental Psychology: Human Learning and Memory, 5, 160–169.

Rouder, J. N., Speckman, P. L., Sun, D., Morey, R. D., & Iverson, G. (2009). Bayesian t tests for accepting and rejecting the null hypothesis. Psychonomic bulletin & review, 16(2), 225–237.

Shankar, K. H., Singh, I., & Howard, M. W. (2016). Neural mechanism to simulate a scale-invariant future. Neural Computation.

Wetzels, R., & Wagenmakers, E.-J. (2012). A default bayesian hypothesis test for correlations and partial correlations. Psychonomic bulletin & review, 19(6), 1057–1064.

Yntema, D. B., & Trask, F. P. (1963). Recall as a search process. Journal of Verbal Learning and Verbal Behavior, 2, 65–74.

